# Epigenetic inhibitors sensitize DLBCL cells to rituximab and doxorubicin

**DOI:** 10.1101/538199

**Authors:** Chiara Facciotto, Julia Casado, Laura Turunen, Suvi-Katri Leivonen, Manuela Tumiati, Ville Rantanen, Liisa Kauppi, Rainer Lehtonen, Sirpa Leppä, Krister Wennerberg, Sampsa Hautaniemi

## Abstract

Primary therapy for diffuse large B-cell lymphoma (DLBCL) is an immunochemotherapy regimen comprising rituximab, cyclophosphamide, doxorubicin, vincristine, and prednisone (R-CHOP). While R-CHOP cures 60% of the patients with DLBCL, those who do not respond or relapse have dismal prognosis, and effective treatment strategies are needed. Due to the plastic nature of the epigenome and its fundamental role in regulating cell identity, we hypothesized that reprogramming the epigenome can overcome resistance to R-CHOP. We developed a novel drug screening protocol to identify epigenetic modifiers that sensitize DLBCL cell lines to immunochemotherapy. Of the herein tested 60 epigenetic compounds, we identified several histone deacetylase (HDAC) and histone methyltransferase (HMT) inhibitors that acted synergistically with immunochemotherapy. We show that sensitization through HDAC and HMT inhibitors is achieved by dysregulating homologous recombination, a central DNA repair pathway, as well as by disrupting the cell cycle and affecting the apoptotic pathway. Epigenetic inhibitors are well-tolerated, which together with our findings support their use in combination with immunochemotherapy in patients with primary refractory and relapsed DLBCL.

## Introduction

Diffuse large B-cell lymphoma (DLBCL) is the most common aggressive lymphoid cancer. The standard of care therapy given to previously untreated DLBCL patients of all ages and subtypes is an immunochemotherapy combination consisting of rituximab, cyclophosphamide, doxorubicin, vincristine, and prednisone (R-CHOP) (*1*), which cures approximately 60% of the DLBCL patients (*1*). Even though several genes, such as TP53, STAT3/6, CDKN2A, and EZH2, have been suggested to confer resistance to R-CHOP (*2, 3*), there is no clinically effective treatment available for R-CHOP resistant patients. Given patients who do not respond to R-CHOP or relapse after primary therapy have dismal prognosis, novel strategies to overcome R-CHOP resistance are needed.

The epigenome dynamically regulates gene expression and is a major player in defining cell identity (*4*). Thus, disrupted epigenome is suggested to cause treatment failure (*5*) and studies have started to investigate how epigenetic reprogramming can revert the resistant phenotype (*6, 7*). Herein we comprehensively assessed the ability of epigenetic inhibitors to overcome treatment resistance in DLBCL. Our approach is based on a novel, high-throughput drug screening protocol that enables non-simultaneous administration of multiple compounds over a period spanning several days. This solves the issues of testing only one or few compounds for a short period of time, which have been major limitations in earlier studies.

We systematically screened 60 epigenetic inhibitors, which allowed to identify effective and clinically usable options to sensitize DLBCL cells to rituximab and doxorubicin, the key compounds in the R-CHOP regimen. Rituximab targets the B-cell surface protein CD20 and its addition to CHOP increased the 5-year overall survival by ~10% (*8–12*), whereas doxorubicin, an anthracycline that induces DNA damage by inhibiting topoisomerase II, increased of 20% the 10-year overall survival (*13, 14*). The epigenetic inhibitors used in the screening target all main classes of epigenetic enzymes, comprising DNA methyltransferases (DNMTs), histone methyltransferases (HMTs), histone acetyltransferases (HATs), histone demethylases (HDMs), histone deacetylases (HDACs), and bromodomains (BRDs).

We discovered that HDAC and HMT inhibitors are particularly effective and thus promising candidates to combination treatment of refractory DLBCL patients. To decipher mechanisms and biomarkers related to epigenetic drug effectiveness, we generated exome and transcriptome sequencing data from DLBCL cell lines before and after treatment. To facilitate exploiting our data and results, we developed an interactive result explorer tool, available at http://app.anduril.org/DLBCL_DSRT.

## Results

### High-throughput multi-step drug combination screening

To systematically investigate the reprogramming ability of multiple epigenetic compounds, we designed the high-throughput screening protocols shown in Figures 1 and S1. Briefly, automated liquid handling allows pretreatment of suspension cells with epigenetic inhibitors followed by exposure to rituximab and doxorubicin. The protocol comprises three main steps. First, cells are seeded on two replicate sets of microplates with previously administered reprogramming compounds. A 10,000-fold concentration range is used to test each epigenetic inhibitor, in order to determine the optimal dose inducing sensitization. Second, cells are incubated with the compounds for either 1 or 3 days (pilot experiment, Fig. S1), or for 9 days using on-plate passaging protocol (Fig. 1). This allows estimation of the time needed by each compound to induce cellular reprogramming. The 9-day pretreatment is too long for cells to survive without fresh media, so we developed a protocol for on-plate cell passaging. After pretreatment, one plate set is treated with rituximab and doxorubicin while keeping another pretreated plate set as control. Third, we measured cell viability to estimate the sensitization induced through epigenetic reprogramming. Protocol details are stated in Materials and Methods.

**Figure 1:**
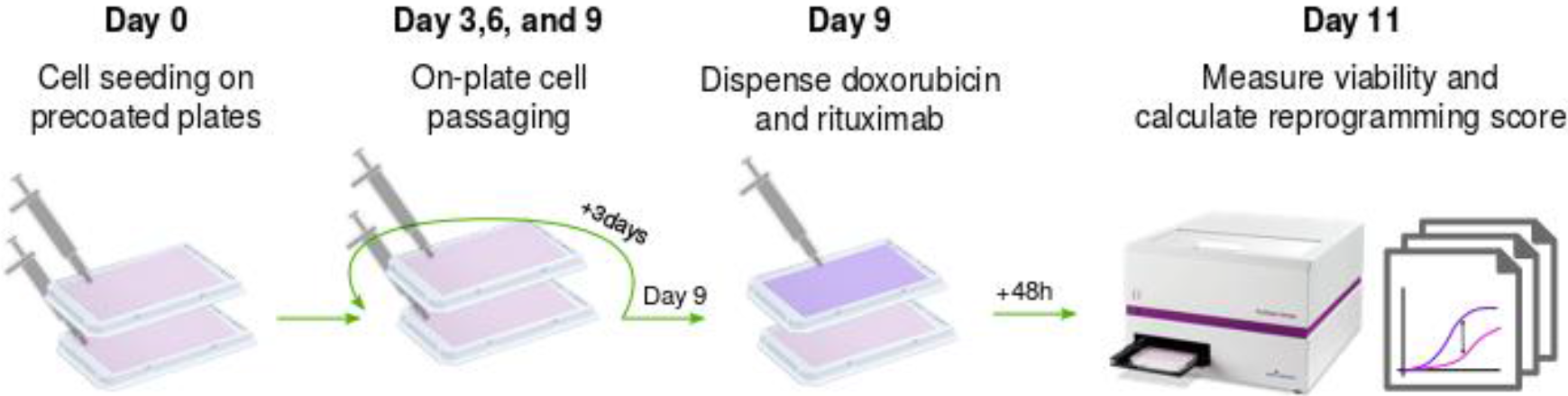
Pretreatment screening protocol designed to simultaneously test the reprogramming activity of 60 epigenetic inhibitors. Cells are seeded on microplates precoated with the pretreatment compounds at five different concentrations. Cells are then passaged in the plate every third day using automated liquid handling. After 9 days of pretreatment, cells are treated with a fixed concentration of doxorubicin and rituximab to compare the activity of the pretreatment alone (pink dose-response curve) vs. the activity in combination with the standard treatment (purple dose-response curve).

### Epigenetic inhibitors sensitize DLBCL cell lines to doxorubicin and rituximab

We applied our novel high-throughput assay to investigate if epigenetic inhibitors could be used to sensitize four DLBCL cell lines (Oci-Ly-3, Riva-I, Su-Dhl-4 and Oci-Ly-19) with varying sensitivity to the combination rituximab and doxorubicin (shown in Figure S2).

We first conducted a pilot screening using short pretreatment times up to three days (described in the Supplementary File and results summarized in Figure S3). This pilot experiment indicated that increasing the length of the pretreatment window enhances the reprogramming ability of epigenetic inhibitors. Thus, we designed our main screening increasing the duration of the pretreatment up to nine days.

We tested 60 inhibitors targeting DNMT (*n* = 7), HDAC (*n* = 21), HAT (*n* = 1), HMT (*n* = 15), HDM (*n* = 3), and BRD (*n* = 13) (Table S1). We measured the sensitization achieved by each compound using a reprogramming score, *i.e.* the maximum difference in cell viability between the effect of the epigenetic inhibitor alone and the effect of the epigenetic inhibitor followed by administration of rituximab and doxorubicin (see Material and Methods).

With the 9-day reprogramming, HDAC inhibitors sensitized all cell lines (Fig. 2C). Furthermore, BRD and HMT inhibitors induced sensitization in three of the four cell lines. Oci-Ly-3 cell line was the most responsive, with 20 out of the 60 epigenetic inhibitors able to sensitize it to doxorubicin and rituximab. Oci-Ly-19 and Su-Dhl-4 cells were sensitized by nine and 10 inhibitors, respectively. Riva-I was the most resistant cell line, successfully reprogrammed only by three inhibitors. The fact that different cell lines responded to different inhibitors was not surprising since compounds such as HDAC inhibitors are known to have different efficacy depending on cancer type and dosage (*15*). The optimal concentration at which each compound induced reprogramming was always lower than the concentration at which the same compound would induce cytotoxicity. Dose response curves are available in the Results Explorer website (http://app.anduril.org/DLBCL_DSRT). A user guide on how to browse data displayed in the results explorer is available in the supplementary materials.

**Figure 2:**
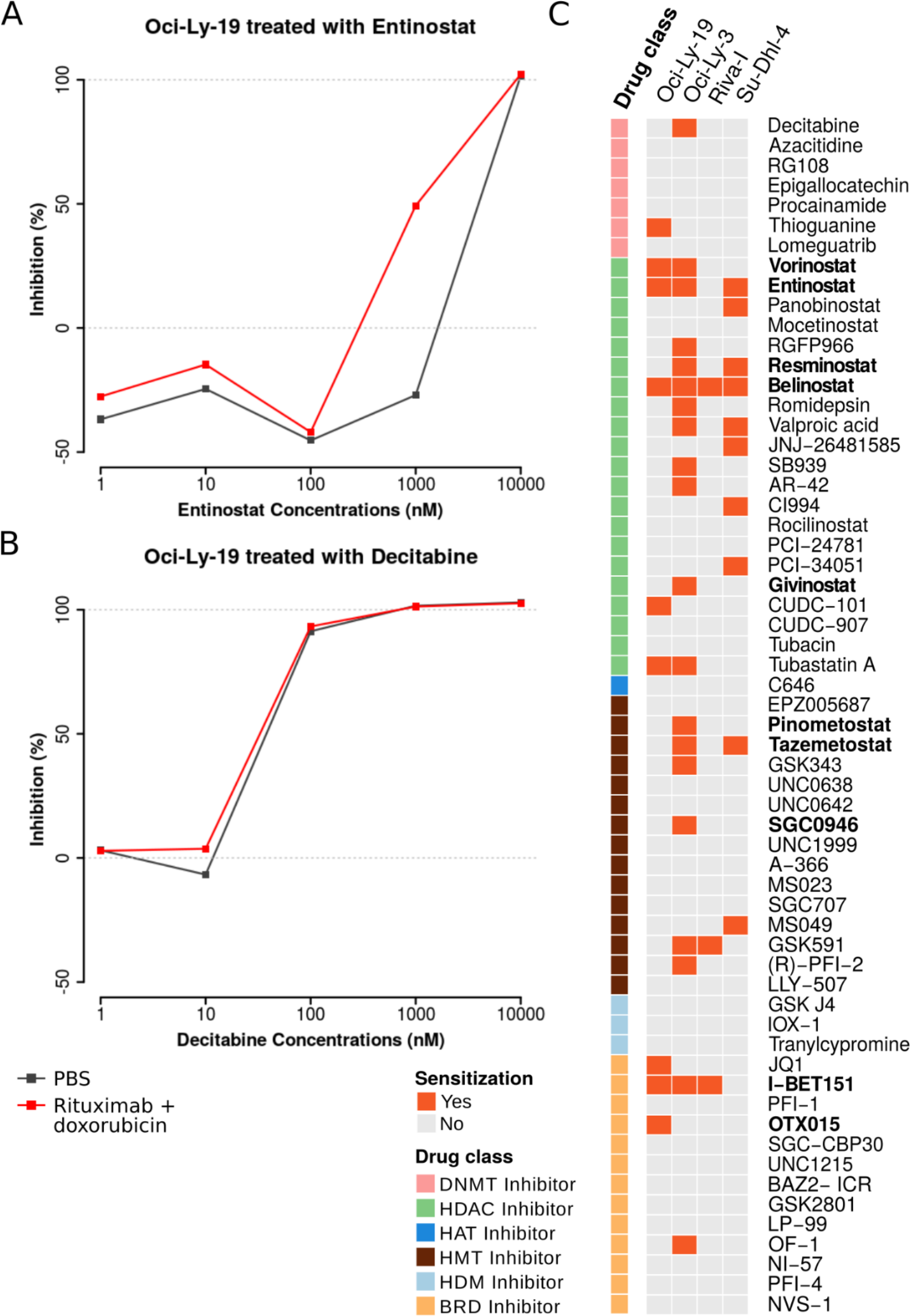
(A) Examples of a compound that induced sensitization to rituximab and doxorubicin vs. (B) a compound that does not sensitize but has a cytotoxic effect. All dose response curves are available in the result explorer. (C) Summary of reprogramming screening hits. Reprogramming scores above a threshold of 30% (see Materials and Methods) and whose dose-response curve passed quality inspection are considered as hits and marked in orange. Ten compounds, marked in bold, were selected for the synergy assay based on their reprogramming potential and mechanisms of action.

### Epigenetic reprogramming acts synergistically with rituximab and doxorubicin

We conducted a drug synergy assay to validate the reprogramming effect of the 10 most potent inhibitors. Compounds with three or more hits, *i.e.*, belinostat, entinostat, and I-BET151, were tested for synergy in all four cell lines, whereas the other compounds were administered only to those cell lines they reprogrammed in the screen. This validation assay followed the same design as the one shown in Figure 1, but varying concentrations of the epigenetic inhibitors, as well as of rituximab and doxorubicin, were now used (see Materials and Methods). Compound concentrations are available in Table S1.

Synergy scores for the 10 inhibitors are shown in Table 1. Scores close to zero indicate that the killing effect of the inhibitors is independent from the killing effect of doxorubicin and rituximab, whereas high scores indicate a synergistic effect (*16*). None of the compounds showed high negative scores, indicating there were no antagonistic effects. The highest synergy scores were observed in compounds targeting either HDACs or HMTs (vorinostat, entinostat, resminostat, belinostat, pinometostat, tazemetostat and SGC0946). Inhibition of BRDs showed lower synergy. The most potent sensitization effects were induced by the HDAC inhibitor entinostat and the HMT inhibitor tazemetostat.

**Table 1:**
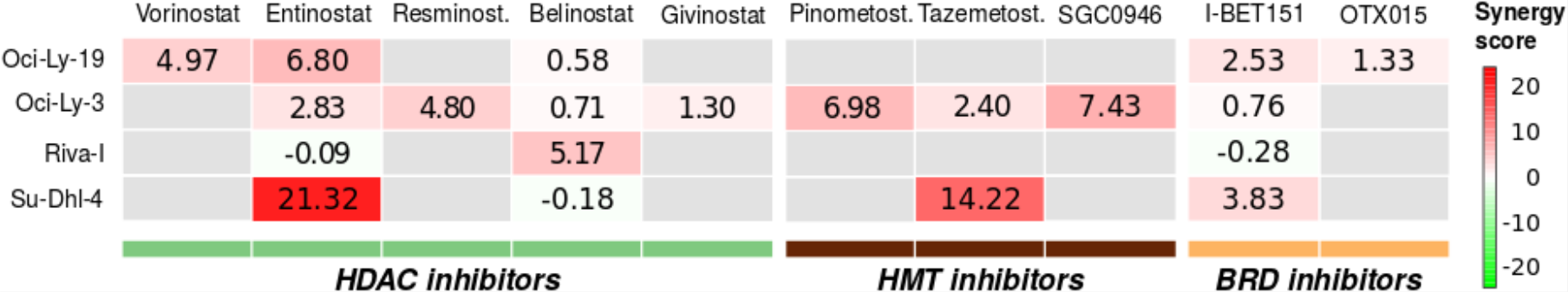
Synergy scores of the top candidate pretreatments after 9-day pretreatment. The figure shows the median scores of three replicate experiments for each measurement where higher score (red) represents synergy with doxorubicin and rituximab, and lower score (green) antagonism. Grey boxes represent untested combinations.

This validation experiment confirmed the findings in the original screen: Oci-Ly-3 cells were the most responsive to reprogramming, and HDAC and HMT inhibitors sensitized them to rituximab and doxorubicin. Su-Dhl-4 and Oci-Ly-19 cells responded to more than one synergistic inhibitor, whereas belinostat was the only compound able to synergistically reprogram Riva-I cells. The synergy plots of the validation experiment are available through the Results Explorer website.

### Epigenetic sensitization to doxorubicin is achieved through reprogramming of DNA repair mechanisms

Since enhanced DNA repair has been suggested to be the key mechanism in doxorubicin resistance (*17*), we conducted an immunofluorescence assay to investigate whether the sensitization effect of HDAC inhibitors is due to impaired repair mechanisms. For this experiment, we selected entinostat, tazemetostat, belinostat, and vorinostat, as they showed high synergy with doxorubicin and rituximab. Of note, these compounds are also clinically relevant as belinostat and vorinostat are already FDA approved for the treatment of patients with relapsed or refractory peripheral T-cell lymphoma and cutaneous T-cell lymphoma, while entinostat and tazemetostat have received FDA “Breakthrough” and “Fast Track” designations.

We used an immunofluorescence assay to measure doxorubicin-induced DNA damage in the form of double strand breaks (DSBs, detected as γH2Ax foci), efficiency of DNA repair via homologous recombination (HR, detected as RAD51 foci) and non-homologous end joining (NHEJ, detected 53BP1 foci), and apoptosis (detected as cleaved-Casp3). Cells treated with HDAC inhibitors (entinostat, vorinostat, belinostat) showed reduced RAD51 focus formation (Figure S4), suggesting impaired HR. NHEJ was upregulated in the HDAC inhibitor treated cells, which was expected as NHEJ is often seen as a compensatory effect for impaired HR.

These results support the hypothesis that HDAC inhibitor sensitization occurs by impairing HR repair as shown in the example reported in Figure 3. Here, entinostat alone does not affect the number of cells positive for DSBs, apoptosis, or HR, compared to the untreated control. However, the response to doxorubicin was strikingly different in cells treated with entinostat compared to untreated cells. The control cells were able to repair DNA damage due to high activity of the HR pathway (green bar) and thus avoid apoptosis.

**Figure 3:**
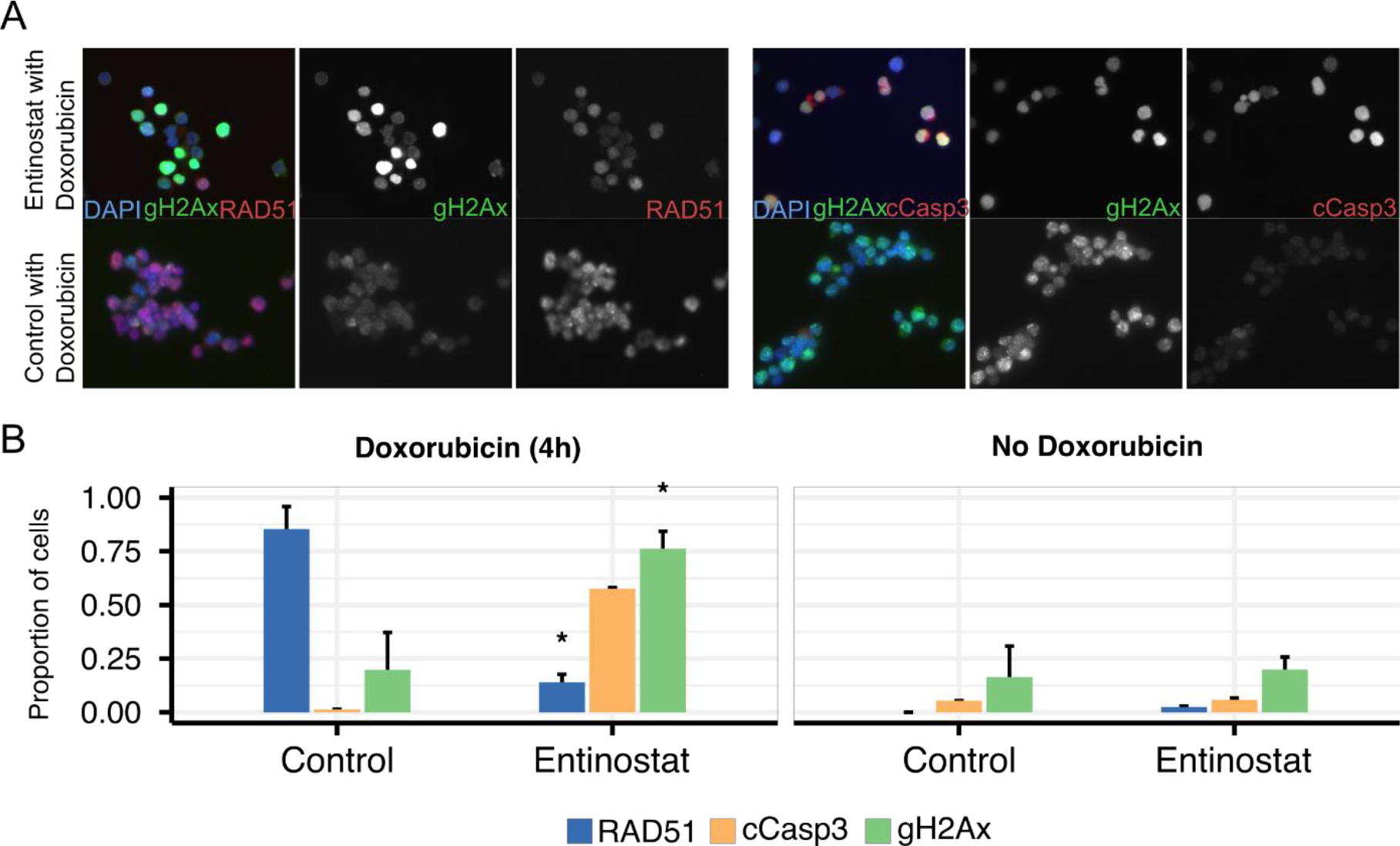
Effect of doxorubicin treatment in Oci-Ly-19 after entinostat treatment on DNA repair mechanisms quantified by immunofluorescence assay. (A) Immunofluorescence images of doxorubicin treated cells after entinostat treatment (above) and untreated (below). Composite image includes also DAPI (blue). (B) Quantification of proportion of cells positive for each marker in each image. The markers shown quantify apoptosis (cCasp3), DNA damage (gH2Ax) and homologous recombination (RAD51) (see methods). The reprogramming effect caused by entinostat under DNA damaging conditions caused by doxorubicin is shown on the left panel, while the right panel shows the lack of effect in the absence of doxorubicin.

### Genomic profiling of DLBCL cell lines

We performed whole exome sequencing (WES) to identify somatic mutations and known germline single nucleotide polymorphisms (SNPs) that could potentially contribute to drug response or synergistic effects of epigenetic sensitizing compounds used in combination with doxorubicin and rituximab.

We used information from public databases and features derived from the WES data to classify variants as likely somatic mutations and likely germline variants (see Materials and Methods). Table S2 sheet “Annotated subset” contains detailed information about manually curated and annotated somatic point mutations (*n* = 14) and germline polymorphisms (*n* = 7) with supporting functional or clinical relevance (discussed in the next paragraph), while the “Filtered variants” sheet lists all 288 variants passing our filtering criteria, of which 268 were classified as somatic and 20 as germline origin (see Materials and Methods). Both somatic and germline variants can be browsed in the Results Explorer website using preloaded (Table S3) or custom gene sets.

At least one potentially functional mutation (Tables 2 and S2) in genes encoding epigenetic enzymes targeted in this study or in genes potentially contributing to response to the epigenetic inhibitory drugs reported in other studies were found in each cell line. A truncating mutation (p.R1322X) in *CREBBP*, which impairs histone acetylation and transcriptional regulation of its targets (*18*) was shared in Riva-I (VAF 0.24) and Oci-Ly-19 (as a subclonal mutation, VAF 07). Truncation of *CREBBP* is acquired in relapse in Acute Lymphocytic Leukemia (ALL) (*18*), further supporting its important role in mediating chemotherapy resistance in lymphoid malignancies. Additionally, Riva-I cells harbor a truncating mutation in the *ARLD1A* gene (p.Q474X), recently reported to encode for a critical transcription factor in the absence of HDAC6 (*19*). ARID1A is the most commonly mutated and functionally disrupted component of the tumor suppressor chromatin remodeling SWI/SNF complex and thereby has been reported to act as an important epigenetic modulator (*20*) and contributor to genetic and genomic instability and response to DNA damaging agents (*21*). Other genes potentially affecting response to the epigenetic inhibitors included BCL6 (epigenetic regulation) harboring a somatic missense mutation in Oci-Ly-3, and STAG2 which contains a truncating substitution, resulting in consistently reduced RNA expression, in Riva-I. STAG2 is a central member of the cohesin complex and if inactivated by such mutations causes genome instability and aneuploidy (*22*). Knock-down of STAG2 is suggested to sensitize pancreatic ductal carcinoma to chemotherapy (*23*).

**Table 2:** Summary of genes carrying functionally annotated somatic mutations and germline polymorphisms* shown in Table S2 sheet “Annotated subset”. These genes have been previously reported as relevant drug targets, epigenetic enzymes, or belong to pathways identified in our analysis.

| Gene Classification |  | Oci-Ly-19 | Oci-Ly-3 | Riva-I | Su-Dhl-4 |
| --- | --- | --- | --- | --- | --- |
| Drug targets or reported association | **Doxorubicin** | *DPYD\**,<br>*NQO1\** | - | *AKT1*,<br>*TP53* | *DPYD\**,<br>*FGFR4\**,<br>*NQO1\**,<br>*TP53* |
|  | **Rituximab** | *DPYD\** | - | *CREBBP* | *DPYD\** |
|  | **Tazemetostat** | - | - | - | *EZH2* |
|  | **Histone deacetylase inhibitors (HDACi)** | - | *MYD88* | *TP53* | *EZH2*,<br>*TP53* |
| Epigenetic | **Histone acetyltransferases (HATs)** | - | - | *CREBBP* | - |
|  | **Histone methyltransferases (HMTs)** | - | - | - | *EZH2* |
|  | **Epigenetic regulation** | *DPYD\** | - | - | *DPYD\** |
| Chemoresponse associated cell mechanisms | **Cell cycle** | - | - | *CREBBP*,<br>*STAG2*,<br>*TP53* | *TP53* |
|  | **DNA repair** | *XRCC3\** | *XRCC3\** | *ERCC4\**,<br>*TP53* | *TP53*,<br>*XRCC3\** |
|  | **Transcription factors** | *CIC* | *BCL6* | *ARID1A*,<br>*CIC* | *RCOR1* |

To characterize the relevance of the identified variants, we used functional prediction and database annotations (described in Materials and Methods) to identify genes targeted by the drugs included in our analysis (listed in Table S4) and to investigate whether such genes harbored mutations (reported in Table 2 and Table S2).

We detected likely pathogenic mutations in doxorubicin related genes (*24*) including *TP53*, *AKT1*, and *EZH2*, and targets of vorinostat *TP53*, *EZH2* and MYD88 (VAF=1 indicates loss of heterozygosity). Tazemetostat also acts through inhibition of *EZH2*, and the specific missense mutation p.Y590S found in Su-Dhl-4 (the tazemetostat responsive cell line) has been reported to increase sensitivity to EZH2 inhibitors (*25*). Notably, the majority of these genes are among the most recurrently altered drivers in DLBCL (*MYD88* in 18%, *CREBBP*, *ARLD1A*, and *TP53* in 10%, and *EZH2* in 6% of patients) (*26*).

Genes somatically mutated or carrying functionally relevant germline polymorphisms in pathways relevant for drug response in our study are summarized in Table 2. Among the genes involved in the repair of DNA damage, *TP53* harbored mutations (p.E162X present in Riva-I cells, and p.R141C present in Su-Dhl-4 cells with VAF 1) and a functionally unconfirmed subclonal *TP53* mutation p.K93R/p.K132R was detected in Riva-I (VAF 0.23) and Oci-Ly-19 (VAF 0.17) cells, while no *TP53* mutations were found in Oci-Ly-3 cells. CIC, a novel transcriptional target of mutant *TP53* and a negative tumor prognostic marker (*27*), was also mutated in Riva-I and Oci-Ly-19 cells. *XRCC3 (28*) and *ERCC4 (29*) are both involved in HR repair and the latter gene also in base excision repair (BER), while *BCL6* is a target of AICDA driven hypermutability process (*30*). All the cell lines except Riva-I displayed a germline risk allele (rs861539) located in the conserved RAD51 domain of *XRCC3*, which has been previously associated to decreased DNA repair capacity in combination with variants in other HR genes (*28*). The G-allele at rs1800124 in *ERCC4* detected in Riva-I has been linked to weaker DNA-protein binding resulting in decreased DNA repair (*31*). Other genes with known germline polymorphisms associated with response to DNA damaging agents commonly used or clinically tested in DLBCL include *NQO1* (rs1800566, linked to response to alkylating agents such as cyclophosphamide, ifosfamide, and platinum compounds), *FGFR4* (rs351855, related to doxorubicin, cyclophosphamide, and fluorouracil response), and *DPYD* (rs1801160, associated to response and rs2297595 to toxicity in to 5-fluorouracil treated patients). Notably, *DPYD* is directly regulated by *EZH2* and repression of DPYD through this mechanism promotes resistance and predicts poor survival in 5-fluorouracil treated patients (*32*).

### Analysis of transcriptome identifies disruption of DNA repair, DNA replication, cell cycle and apoptosis pathways aspotential mechanisms behind epigenetic sensitization

To further characterize the molecular mechanisms affected by epigenetic sensitization, we performed RNA-seq of the four cell lines before and after treating them with belinostat, entinostat, vorinostat, and tazemetostat (Figure S5A). Differentially expressed genes (DEGs) between treated and untreated cells are shown in Figure S5B-E, and are also available in the Result Explorer website.

We used DEGs from each successfully reprogrammed combination and performed pathway enrichment analysis to explore the mechanisms affected by epigenetic reprogramming (Table S5). All reprogrammed combinations showed changes in the mechanisms related to immune response. This was expected since DLBCL originates from B-cells, which produce antibodies in the adaptive immune system (*33*). Our pathway analyses further revealed the major histocompatibility complex (MHC) as one of the most affected pathways for HDAC inhibitors, which is in line with a recent study (*15*).

DNA damage and repair pathways were dysregulated in Su-Dhl-4 and Oci-Ly-19 cells treated with entinostat, as well as in Su-Dhl-4 cells treated with tazemetostat. When comparing the untreated and treated conditions for all successfully sensitized cell line and inhibitor combinations, we identified DEGs belonging to the HR, NHEJ and other DNA repair pathways (Table S6). While DEGs belonging to the NHEJ pathways were identified only in Su-Dhl-4 cells treated with entinostat, HR genes were differentially expressed both in Su-Dhl-4 cells treated with entinostat and tazemetostat, as well as in Oci-Ly-19 cells treated with entinostat and vorinostat. In particular, these combinations showed downregulation of both *XRCC2* and *POLQ* expression.XRCC2 is essential for the proper functioning of the HR pathway (*34*), and knockdown of *POLQ* in HR-deficient tumors enhances cell death (*35*). Thus, decreased expression of both genes may contribute to the sensitization of doxorubicin-resistant DLBCLs. Additionally, Su-Dhl-4 cells treated with entinostat or tazemetostat also showed downregulation of *RAD51*, *RAD54L*, *BRCA2*, and three Fanconi anemia genes (*FANCA*, *FANCB*, and *FANCM*). *BRCA1* expression was suppressed in Su-Dhl-4 cells in response to entinostat. In Riva-I cells, belinostat-induced differential expression did not have an impact on the expression of any of the HR genes, making it the only cell line reprogrammed by a HDAC inhibition, but not showing transcriptional changes in DNA repair.

Disruption of cell cycle and DNA replication were also mechanisms identified in several sensitized combinations (*i.e.*, Su-Dhl-4 and Oci-Ly-19 cells treated with entinostat, as well as Su-Dhl-4 cells treated with tazemetostat). In particular, treatment of Su-Dhl-4 cells with entinostat led to upregulation of *CDKN1A*, an HDAC inhibitor mechanism previously suggested to induce cell cycle arrest (*15*).

Other pathways identified in our analysis included cell adhesion (altered in all sensitized combinations except Riva-I cells treated with belinostat) and TGFP signaling (disrupted in Oci-Ly-19 cells treated with either entinostat or vorinostat, and in Su-Dhl-4 cells treated with entinostat). Death receptors and ligands belonging to the TNF and TNF-receptor superfamilies were differentially expressed in all sensitized combinations. Indeed, HDAC inhibitors have been demonstrated to affect apoptosis through dysregulation of such protein families (*15*). Pathway analysis also showed disruption of the apoptotic pathway in Riva-I cells treated with belinostat.

### Increased CD20 expression does not sensitize all cell lines to rituximab

Next, we investigated whether rituximab sensitization in response to epigenetic modifiers was a result of increased expression of the *MS4A1* gene encoding for CD20. None of the four tested epigenetic compounds upregulated *MS4A1* gene expression in any of the cell lines. However, *CD40*, which is a key effector of CD20 on B cells (*36*), was inhibited in Oci-Ly-19 cells in response to vorinostat or tazemetostat.

We also examined if sensitization to rituximab could be due to increased expression of CD20 on the lymphoma cell surface. Two compounds shown to support CD20 transport to the cell membrane were tested using our high-throughput screening (Figure S6). Rifampicin, an antibiotic shown to restore efficacy of anti-CD20 antibodies (*37*), was successful only on Oci-Ly-19 cell line. Suramin, a small molecule that inhibits CD40, sensitized the rest of the cell lines. However, since this sensitization was observed at different pretreatment time points for different cell lines, we cannot conclude that upregulation of CD20 expression is the key factor for sensitization to rituximab.

When comparing the expression of rituximab related genes (*24*) in the untreated cells, we observed that Oci-Ly-3 cells presented a unique profile compared with the other cell lines. Specifically, Oci-Ly-3 was the only cell line showing overexpression of *CXCL13*, a B cell-attracting chemokine (*38*), and downregulation of *CD27*, aproapoptotic gene previously shown to link HDACs activation and cell cycle arrest (*39*).

## Discussion

Finding effective treatment options for relapsed and refractory malignancies is a major challenge in cancer therapy. We developed a high-throughput experimental protocol that allowed the identification of epigenetic inhibitors able to (re)sensitize cancer cells to standard therapeutic agents. This approach is a major step towards finding clinically useful and effective drug combinations. Here, we demonstrated how the protocol could assess the sensitization power of 60 epigenetic modifiers in DLBCL. Moreover, the customizable plate layout and the ability to vary experiment duration by on-plate cell passaging makes this protocol suitable for testing other drug combinations with multi-step delivery.

Due to the plasticity of the epigenome, epigenetic inhibitors are a particularly interesting class of drugs to sensitize cancer cells to standard therapeutic options (*40*). Our results indicate that most of the epigenetic inhibitors require several days to effectively induce reprogramming. Only a few compounds, mainly HDAC inhibitors, were able to sensitize cell lines within one and three days of pretreatment. It is not surprising to observe that certain classes of epigenetic inhibitors are not fast-acting, especially when considering their mechanisms of action. For instance, DNMT inhibitors are expected to be slow acting, since passive demethylation requires several cell cycles. However, pairing nine days pretreatment time with multiple doses of the inhibitors sensitized all cell lines to doxorubicin and rituximab, the key compounds of the R-CHOP regimen. This suggests that epigenetic reprogramming is effective across all molecular subtypes.

Our results show that epigenetic inhibitors should be administered in a reprogramming mode, *i.e.*, several doses and days before chemotherapy, rather than simultaneously with chemotherapy. Importantly, this study demonstrates that epigenetic drugs induced sensitization at much lower doses than those required for cytotoxicity. This suggests that when epigenetic drugs are used to sensitize rather than kill cancer cells, they are likely to cause less severe side-effects.

HDAC inhibitors, most notably belinostat, entinostat, vorinostat, and resminostat, as well as HMT inhibitors such as tazemetostat, pinometostat, and SGC0946, were the most potent epigenetic drugs to sensitize cancer cells to doxorubicin and rituximab. Both classes of inhibitors are reported to be well-tolerated in clinical trials (*15, 41, 42*), hence their reprogramming potential should be further investigated *in vivo* to identify the correct time and dose for epigenetic sensitization.

When looking into the mechanisms behind epigenetic reprogramming, dysregulation of DNA repair pathways (especially HR), disruption of cell cycle and effects on apoptosis were among the main ones identified in our study. This corroborates previous observations on the mode of action of HDAC and HMT inhibitors (*15, 43*).

Resistance to DNA damaging agents, such as doxorubicin (*44*), has often been associated with upregulation of HR-mediated DNA repair. Our results show that belinostat, entinostat, vorinostat and tazemetostat can sensitize DLBCL cells via disrupting the HR pathway. Further, sensitization by these inhibitors affected the expression several DSB repair genes, such as *XRCC2* and *POLQ*, both downregulated in sensitized combinations (except Riva-I cells treated with belinostat). These findings argue that epigenetic inhibitors could also revert resistance to a wider class of DNA damaging agents, such as platinum-based regimens and radiotherapy. Belinostat-treated Riva-I cells showed activation of the HR pathway only in the immunofluorescence assay, while no transcriptomic changes were found in the DNA repair genes. However, *ARID1A* mutation in Riva-I cells might explain its sensitivity to pan-HDAC inhibitors, such as belinostat (*19*). In addition, apoptosis mechanisms were severely disrupted in this combination, suggesting that HR dysregulation may not be the only pathway affected by belinostat-mediated sensitization.

Even though we found a clear link between epigenetic modifiers and sensitization to doxorubicin, we were unable to find the same for rituximab. The major reasons are the undefined mechanisms of action of CD20 *in vitro* and the absence of known regulatory pathways modulating this protein. However, the smaller impact of rituximab to therapy response compared to that of DNA damaging agents (*e.g.*, doxorubicin) corroborates the weak link between rituximab and epigenetic reprogramming.

Through the genomic characterization of the DLBCL cell lines we were able to highlight several mutations with functional or clinical relevance. The most striking result was the identification of a missense mutation on gene *EZH2* in the GCB cell line Su-Dhl-4, showing the highest response to EZH2-inhibitor tazemetostat. This EZH2 inhibitor has the highest efficacy in *EZH2*-mutated DLBCL patients belonging to the GCB subtype (*45*), and the mutation found in Su-Dhl-4 cells has been reported as a potential biomarker for response to EZH2 inhibitors (*46*). Moreover, tazemetostat treatment significantly deregulated the expression of cell cycle genes in this cell line, potentially causing cell cycle arrest, a mechanism suggested by Knutson *et al. (43*). Thus, our results are in line with the earlier clinical findings and support the predictive value of mutations in the *EZH2* gene as biomarkers for the selection of DLBCLs that could be sensitized by tazemetostat. Altogether, these results suggest that R-CHOP resistant DLBCL patients with *EZH2* mutations could benefit from tazemetostat pretreatment followed by R-CHOP re-challenge.

We are aware of our study limitations. First, when we created the compound library, only fewHDM and HAT inhibitors were available. In our experiments, these compound classes were not able to induce reprogramming, but testing a more extensive collection of inhibitors might identify mechanisms able to induce reprogramming by targeting HDM and HAT enzymes. Second, even though all DLBCL cell-of-origin subtypes were represented in our study, the number of cell lines included was relatively small. However, since our protocol is high-throughput, this kind of experiment can be scaled up in future studies. Third, the impact of tumor microenvironment, which is an important mediator of rituximab response in B-cell lymphomas could not be tested in this model. Even though our results suggest that many epigenetic inhibitors can be useful in a clinical setting, a more detailed analysis is needed to estimate the optimal dose and treatment duration *in vivo.*

Taken together, our results support the application of epigenetic reprogramming in sensitizing DLBCL to standard immunochemotherapy combinations. Our study is among the few investigating the use of epigenetic drugs as sensitizing agents, instead of using them as mono- or combination therapy, and it is the first to do so in a systematic and high-throughput fashion. Our findings warrant further investigation of epigenetic inhibitors as sensitizing agents to treat patients with primary refractory and relapsed DLBCL.

## Materials and Methods

### Study design

We investigated the sensitizing effect of 60 epigenetic inhibitors (listed in Figure 2 and Table S1) to rituximab and doxorubicin on DLBCL cell lines *in vitro.* We developed and performed a high-throughput assay to screen their sensitizing potential using treatment windows of different length (Figure 1). After validating our findings, we further investigated the effect of epigenetic sensitization on different pathways using a combination of immunofluorescence assays and sequencing technologies.

### Compound collection design

We curated the compound library by manually searching literature and providers for compounds which can be used to identify the epigenetic mechanisms leading to tumor sensitization, comprising both FDA-approved compounds and probes.

Our collection included inhibitors of all the main families of epigenetic enzymes currently under investigation. The full list of compounds is available in Figure 2 and in Table S1. The library for the longest pretreatment time included also 16 compounds (SGC0946, UNC1999, A-366, MS023, SGC707, MS049, GSK591, (R)-PFI-2, LLY-507, BAZ2-ICR, GSK2801, LP-99, OF-1, NI-57, PFI-4, and NVS-1) obtained through an MTA with the Structural Genomics Consortium (SGC). The remaining compounds were obtained from commercial sources by the High Throughput Biomedicine Unit of the Institute for Molecular Medicine Finland (FIMM), University of Helsinki.

### Cell lines

We selected four DLBCL cell lines, representative of all DLBCL subtypes and with varying response to rituximab and doxorubicin (Figure S2). Cell lines were kindly provided by Dybkμr lab (Aalborg University) and they represent all DLBCL molecular subtypes (*47*). Su-Dhl-4 belongs to the GCB subtype (*48*), Oci-Ly-3 and Riva-I to the ABC subtype (*48*), while Oci-Ly-19 is unclassified. All four cell lines express the *MS4A1* gene (the gene encoding the CD20 protein), with Oci-Ly-19 showing the lowest expression according to our transcriptomic data. According to DSMZ cell line bank, Oci-Ly-19 is 4% polyploid which is consistent with our observations suggesting hyperpolyploid nature of this cell line.

### Cell culture conditions

Riva-I, Su-Dhl-4 and Oci-Ly-19 were cultured in RPMI medium with 10% FBS, and Oci-Ly-was cultured in IMDM medium with 20% FBS. Cells were passaged every second day with a ratio of 1:2 (Su-Dhl-4, Oci-Ly-3) or 1:3 (Riva-I, Oci-Ly-19). Cells were kept in an incubator at a constant 37°C and 5% CO_2_. All cell lines were authenticated by STR analyses and tested negative for mycoplasma contamination.

### Screeningprocedure and parameters

Compounds were dissolved in DMSO and added to the assay plates using a Labcyte Echo 550 acoustic dispenser. The highest dose concentration was as advised by the supplier followed by four 10-fold dilutions. Details on each compound’s individual dose can be found in Table S1. Plate layout design included randomized positive (benzethonium chloride, BzCl, Sigma-Aldrich) and negative (DMSO, Sigma-Aldrich) controls. Compound plates were stored under inert nitrogen gas in StoragePods (Roylan Developments) until needed. Cells were seeded using BioTek MultiFlo FX Random Access Dispenser, at 3000 cells/well (Riva-I, Su-Dhl-4, Oci-Ly-19) or 4000 cells/well (Oci-Ly-3) in 25 *μ*L (1 and 3 days of pretreatment time) or 40 *μ*L (9 days pretreatment time). Cell plates were incubated in a Thermo Scientific Cytomat 10C incubator at 37°C and 5% CO_2_. Plates undergoing 9 days of pretreatment had a Labcyte microclime lid to reduce media evaporation during the incubation period. During the longest pretreatment time, cells were passaged every third day (*i.e.* on day 3, 6 and 9) directly in-plate with pretreatment drugs added to the new media. Since all cell lines grew in suspension, plates were spun down before passaging. In-plate passaging was then performed with a Beckman Coulter Biomek FXp pipetting device fitted with a 384 multichannel head. The BioMek FXp protocol included the following steps:

1. Aspirate 20 *μ*L of old media from the culture plate (without touching the cells collected at the bottom of the well) and discard it.
2. Aspirate 20 *μ*L of fresh media from the plate with drugs in media and dispense it on the cells.
3. Resuspend the cells by mixing with 20 *μ*L volume five times.
4. Aspirate 20 *μ*L of old media and cells from the culture plate and discard it.
5. Aspirate 20 *μ*L of fresh media from the plate with drugs in media and dispense it on the cells.

With this procedure roughly 3/4 of the media was exchanged while removing half of the cells from each well.

After pretreatment, half of the plates were treated with a fixed dose of rituximab (MabThera, diluted in PBS. Roche) and doxorubicin (diluted in PBS. Sigma-Aldrich), while the other half received only PBS as control. The concentrations of rituximab and doxorubicin were determined through drug a combination assay and are listed in Table S1. After treatment, cells were incubated for 48 h. Finally, cell viability was measured with Promega CellTiter-Glo reagent and BMGLABTECH FLUOstar Omega plate reader.

### Cell lines response to rituximab and doxorubicin

Rituximab and doxorubicin were selected because they are the main contributors of R-CHOP. While the main mechanism of action of monoclonal antibodies is to reactivate the immune response, rituximab can also induce direct apoptosis (*49, 50*), making it suitable for this *in vitro* study.

The static concentration of the combination of doxorubicin and rituximab was selected by performing a drug combination assay. To emulate the conditions of the cell culture at the moment of treatment in the screening procedure, this assay followed the same protocol without pretreating with epigenetic inhibitors. On treatment day the cells are treated with combinations of seven 10-fold dilutions of doxorubicin and rituximab starting at 1000mM and 10mg/mL respectively. The plate layout included triplicates for each combination dose.

### Microplate reader dataprocessing

All dose response analyses were implemented within the Anduril framework (*51*). Quality control and filtering was done using positive (BzCl, cell killing) and negative (DMSO) controls. This step comprises analyzing mean, standard deviation, coefficient of variation (CV), signal to background ratio, Z’-factor, and strictly standardized mean difference (SSMD) (*52*). Plates with a CV higher than 20% were discarded. Each plate was normalized using the mean of negative controls as 0% inhibition and the mean of positive controls 100%.

### Reprogramming scores

The reprogramming score for a pair, pretreatment and cell line, is a score between 0 and 100 defined as follows:

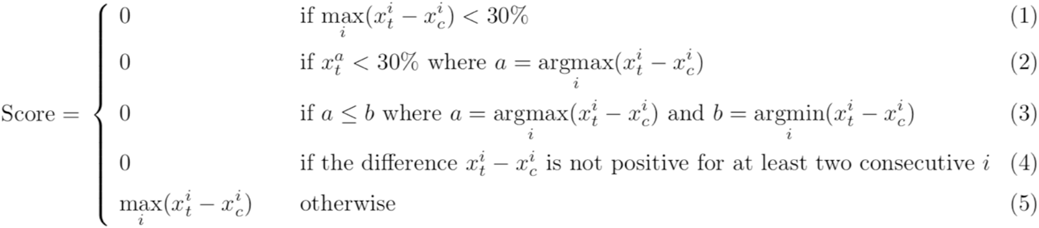

where *i* can be any of the five doses of pretreatment compound tested, and *x_t_* and *x_c_* are the values of normalized inhibition observed in the treatment plate (the plate that was additionally treated with rituximab and doxorubicin after pretreatment time) and the control plate (the plate that only received the pretreatment) respectively. In brief, the score is defined as the highest difference between treatment and control dose-response curves, 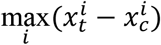, unless it is truncated to zero. This truncation was added to ensure that the scores represent actual reprogramming events and not the effect of outliers. The algorithm returns 0 if (1) the highest difference is not large enough to consider a relevant increase on cytotoxicity, (2) despite showing reprogramming effect the inhibition achieved is low, (3) the dose that achieves the highest difference is not larger than the dose that achieves the minimum difference, (4) the reprogramming effect is not consistent throughout the dose-response curve, e.g. due to outliers. Additionally, all dose-response curves were inspected manually to confirm the quality of the data. The standard method of dose-response curves to a sigmoidal function did not apply to this data due to unexpected effects caused by epigenetic reprogramming, such as enhanced cell growth, therefore a partial function was needed to determine which compounds could effectively induce epigenetic sensitization.

### Synergy assay between epigenetic pretreatment and doxorubicin-rituximab combination therapy

The pairs, epigenetic compound and cell line, to be validated in this assay were selected from the results shown in Figure 2 based on reprogramming scores combined with manual inspection of dose-response curves. The plate design included all combinations of epigenetic pretreatment with rituximab and doxorubicin using 5 concentrations for each compound and 5 concentrations for rituximab and doxorubicin as a combination matrix layout. Three replicates of each matrix were included in different locations in the plate. The dose ranges are detailed in Supplementary Table S1. Synergy scores were calculated with the synergyfinder tool (*53*) applying the zero interaction potency model (ZIP), which compares the change in inhibition of the dose-response curves between individual drugs and their combinations, the final score quantifies the deviation from the expected inhibition in the case of zero interaction (*54*). The scores are available in our Results Explorer (see Data and material availability) under “Synergy Investigation”.

### DNA and RNA sequencing

All cell lines were analyzed by whole exome sequencing (WES) and RNA sequencing before and after pretreatment. DNA and RNA extraction were performed using NucleoSpin Tissue (Macherey-Nagel) and NucleoSpin RNA Plus (Macherey-Nagel) respectively.

WES target enrichment was done with SureSelect Human Exome V5 baits (targeting 50 Mb of exonic regions in 31,522 genes) and 350 bp inset size libraries were constructed with SureSelect XT library kit (Agilent Corporation, CA, USA) according to manufacturer’s protocols.

The amount, concentration and integrity of the RNA samples were estimated with Bioanalyzer 2100 (Agilent) at the Biomedicum Functional Genomics Unit (Helsinki, Finland). Total RNA fragments (140-160 bp) including coding and long non-coding RNAs were sequenced using a strand specific protocol similar to TruSeq Stranded Total RNA (Illumina) including removal of ribosomal RNA with Ribo-Zero™ Magnetic Kit and RNaseH (Illumina) and random hexamer primers.

DNA and RNA samples were sequenced by BGI Genomics Co., Ltd. (Hong Kong) with Illumina HiSeq4000 sequencer and chemistry (Illumina Inc. CA, USA) using a standard paired-end protocol with 100 bp read length. In total, 150 Gb of clean data to reach 50x target coverage in WES and 60 million 100 bp reads for whole transcriptome analyses in RNA-Seq were produced from each sample.

### Whole Exome Sequence analysis

All analyses were performed by custom bioinformatics pipelines within the Anduril framework (*51*) to automate all steps. Quality control was done with FastQC (*55*) and read ends trimmed by trimmomatic (*56*). After discarding low quality reads, and single strand reads, 75% of the reads from each sample were kept. Sequence alignment to reference genome hg19 was done with Burrows-Wheeler Aligner (*57*) and sorted with Picard tools (Broad Institute), 90% of the targets had coverage >10x for all the cell lines (average 33-34x). Variant calling and variant filtering were performed following GATK recommended practices (*58*) and Annovar (*59*). A preliminary filter kept only splicing and exonic variants with VAF higher than 20% in at least one cell line, CADD score higher than 10 and COSMIC (*60*) annotation except in cases where no SNPdb annotation was available. Then all variants within genes with FPKM less than 1 in our RNA-seq data were discarded. Additional filtering steps were performed to identify relevant variants. We defined a variant as somatic if it was present in less than three cell lines and had Minor Allele Frequency (MAF) less than 1% given by Annovar MaxPopFreq. Variants with MAF up to 5% were also considered somatic in entries with over six occurrences reported as somatic in COSMIC. The rest of the variants were assessed as germline. We filtered germline variants based on potential association to drugs tested for synergy with epigenetic inhibitors and if they were supported by public databases or literature. Moreover, variants annotated in ClinVar (*61*) were retained for further evaluation. In general, clinical relevance and drug-gene/drug-variant interactions were annotated using the following data sources: CIVIC (*62*), DGIdb (*63*), DrugBank (*64*), and PharmGKB (*24*). Cancer driver status and drug response association of all protein alterations were predicted using Cancer Genome Interpreter (*65*). Variants of selected genes were further annotated using gene-specific data bases, LOVD3 (*66*) for *BRCA1* and *RAD51*, and IARC for *TP53 (67*).

### RNA-seq analysis

Paired-ended fastq reads were preprocessed using an Anduril pipeline that combines state-of-the-art tools. Read quality before and after trimming was accessed using FastQC (*55*). Adaptor removal and trimming were carried out with trimmomatic (*56*) using the following parameters: headcrop = 10, slidingWindow = 5:20, minQuality = 30, trailing = 30. Since for each sample more than 70% of all reads had quality above 30 and were more than 20 base pairs long after the trimming step, no dataset was discarded. Alignment was performed using two-pass STAR (*68*). Reads were aligned to reference genome hg19 and the software was run using the default STAR parameters. Gene expression was then quantified using eXpress (*69*). Genes not expressed in both the treated or untreated condition (i.e. log2 FPKM < 1) were excluded from the analysis. Normalization and rlog transformation of gene count data was computed with the R package DESeq2 (*70*), and normalized data were used to compute differential expression. Log2 fold change and absolute difference where computed between the untreated and treated pairs for a total of 14 comparisons. Genes with absolute log2 fold change > 1 and absolute difference > 1 were classified as differentially expressed genes (DEGs). Due to the experimental design of this study and the absence of replicates, no p-value could be computed, and the significance of the differential expression could not be estimated. We checked if any of the DEGs in such sets included any DSBs DNA repair genes. We also performed pathway enrichment for the DEGs belonging to one of the five sensitized combinations (Oci-Ly-19 entinostat, Oci-Ly-19 vorinostat, Riva-I belinostat, Su-Dhl-4 entinostat, and Su-Dhl-4 tazemetostat). We repeated this analysis using both all DEGs identified in each combination as well as those subsets of DEGs shared by all cell lines sensitized by each inhibitor but absent in those not sensitized (see sets highlighted in yellow in Figures S5B-E). The R package EnrichR (*71*) was used to perform pathway enrichment using KEGG (*72*), WikiPathways (*73*), and Reactome (*74*) as reference databases. Pathways with at least three genes differentially expressed and with p-value less than 0.05 are listed in Table S4.

### Analysis of DNA damage, DNA repair and apoptosis

Cells were cultured in T-25 flasks and treated with the epigenetic compounds for nine days. Untreated cells were used to assess endogenous levels of DNA damage and apoptosis. To induce DNA damage, cells were then exposed to 1 *μ*M (Su-Dhl-4, Oci-Ly-19, Oci-Ly-3) or 100 *μ*M (Riva-I) of doxorubicin. Cells were collected after 4 and 24 h, centrifuged, washed with PBS, fixed with 2% buffered paraformaldehyde and spread on coated microscope slides for immunostaining. Primary antibodies against yH2Ax (ab22551, Abcam), RAD51 (sc-8349, Santa Cruz Biotechnology) and cleaved caspase 3 (9664, Cell Signaling Technology) were used to detect DNA damage, homologous recombination-mediated DNA repair and apoptosis, respectively, as previously described (*75*). Images were acquired on a Nikon Eclipse-90i epifluorescence microscope. Image segmentation and quantification were performed using a custom script in the Anima framework (*76*), followed by statistical summary in R; mean, standard deviation, standard error, and confidence intervals (Figure S4).

### Results explorer

The results explorer was created using the R package Shiny (77). Plotly (*78*) was used to produce the interactive plots in the “Reprogramming Screening” and “Synergy Investigation” tabs, while the heatmaps in the “Gene Expression” and “Mutations” tabs were created using heatmaply (*79*). The full list of genes included in the preloaded sets are available in Table S3.

## Supplementary Materials

Fig. S1. Experimental procedure for 1 and 3 days of pretreatment.

Fig. S2. Dose response curves of rituximab and doxorubicin for all four cell lines.

Pilot epigenetic screening with pretreatment time up to three days.

Fig. S3. Screening results for 1 and 3 days of pretreatment.

Fig. S4. Results from immunofluorescence assay investigating DNA repair.

Fig. S5. Summary of the sequencing data and of differentially expressed genes.

Fig. S6. Sensitizing effect of CD20-transport modulators.

Table S1. Reprogramming scores, synergy scores and compound concentrations (Excel)

Table S2. DLBCL cell line exome sequencing key somatic mutations and germline polymorphisms with annotations. (Excel)

Table S3. Gene sets preloaded in the result explorer (Excel)

Table S4. Target genes of drugs used in the synergy experiment (Excel)

Table S5. Pathway enrichment results using EnrichR (Excel)

Table S6. DEGs belonging to DNA repair pathways (Excel)

Results Explorer - User Guide

## Acknowledgements

The authors thank Dr. Karen Dybkær from Aalborg University, Denmark, for providing us the cell lines used in this study, Structural Genomics Consortium (SGC) for sharing the 16 compounds tested in the 9-day pretreatment, and IT Center for Science (CSC) for computing resources. The authors also thank Anne Aarnio, Marika Tuukkanen, and Kirsi Jantti for their guidance and technical assistance, Jani Saarela for the helpful discussions about drug screening data analysis.

## Author contributions

C.F. and J.C. contributed equally to the study and wrote the manuscript. C.F. and J.C. conceptualized the study together with S.H., R.L., S.L., K.W., S.-K.L. and L.T. F., J.C., L.T., V.R. andM.T. performed experiments. R.L., S-K.L., S.L. and S.H. supervised the study; L.K., S.L. and S.H. acquired funding. All authors read and edited the paper.

## Data and materials availability

Results from the drug screening assays and from sequencing data analyses can be browsed in our result explorer at http://app.anduril.org/DLBCL_DSRT:

- The “Reprogramming Screening” tab shows the results of the assay with five concentrations of the epigenetic inhibitors and one fixed concentration of rituximab and doxorubicin. Results for each epigenetic compound can be grouped by length of pretreatment or by cell lines for comparisons.
- The “Synergy Investigation” tab shows the results of the 9-days synergy assay with five concentrations of the epigenetic inhibitors and five concentrations of rituximab and doxorubicin.
- The “Gene expression” tab allows to compare the expression of a pre-selected set of genes across cell lines that are untreated or treated with epigenetic compounds, to access how the reprogramming affected the transcriptome. Some gene sets are preloaded, but the user can load new sets for customized analyses.
- The “Mutations” tab allows to visualize which genes carry mutations in the cell lines included in this study. Some gene sets are preloaded, but the user can load new sets for customized analyses.
- The “Immunofluorescence” tab allows to visualize the images obtained through the immunofluorescence assay used to investigate DNA damage and repair.

The raw fastq files from our WES and RNA-Seq experiments are available in SRA (*80*), at the accession number PRJNA517451.

