## Supplementary File for "Epigenetic inhibitors sensitize DLBCL cells to rituximab and doxorubicin"

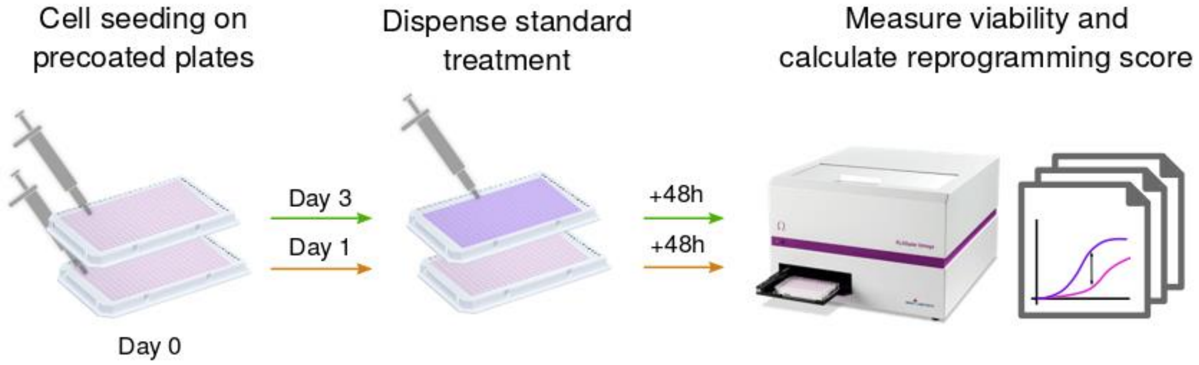


***Supplementary Figure S1:*** *Pretreatment screening protocol designed to simultaneously test the reprogramming activity of 44 epigenetic inhibitors. Lymphoma cells are seeded on microplates precoated with the pretreatment compounds at five different concentrations. After the pretreatment time (1 or 3 days), cells are treated with a fixed concentration of doxorubicin and rituximab in order to compare the activity of the pretreatment alone (pink dose-response curve) vs. the activity in combination with the standard treatment (purple dose-response curve).*

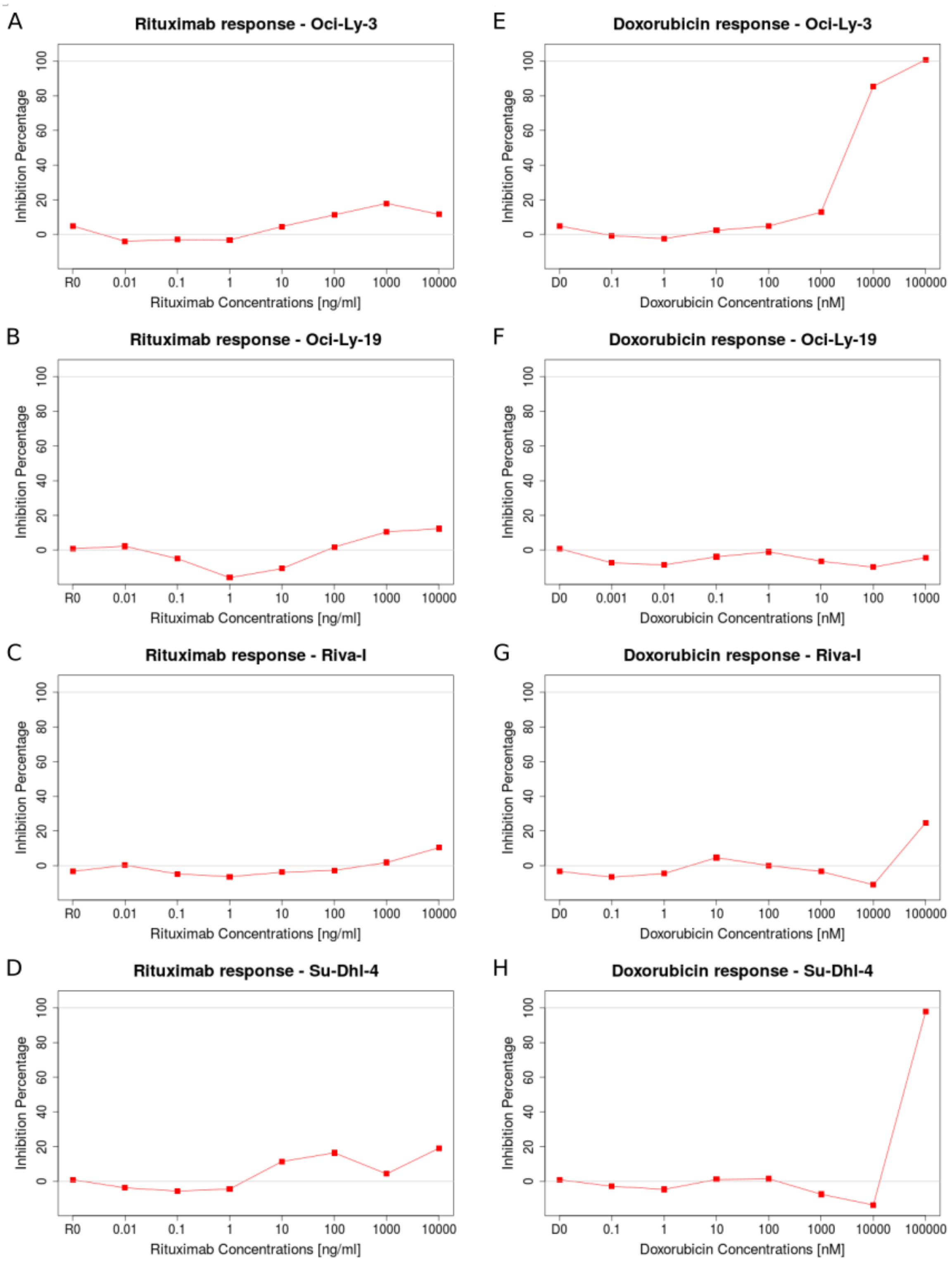


**Supplementary Figure S2:** Dose responses showing the level of rituximab (A-D) and doxorubicin (E-H) resistance of each cell line. R0 and D0 represent measurements of cell viability in the absence of rituximab and doxorubicin respectively.

*Pilot epigenetic screening with pretreatment time up to three days*

When developing our screening protocol, we first conducted a pilot screening (Figure S1) with short pretreatment times (one and three days) to test 44 inhibitors targeting DNMT ($n$ = 7), HDAC ($n$ = 21), HAT ($n$ = 1), HMT ($n$ = 6), HDM ($n$ = 3), and BRD ($n$ = 6) (Table S1). Sensitization was estimated by computing a reprogramming score (Table S1) for each combination of compound and cell line (see Material and Methods). Dose response curves for this pilot experiment can be found in the result explorer, while Figure S3 summarizes which compounds successfully induced sensitization in our cell lines.

Pretreating the cells for one day induced sensitization mainly in Su-Dhl-4 cells. However, most of the potential hits occurred at low doses. Since such effect was lost when increasing the pretreatment time, we decided to further investigate it with a second screening including compounds AR-42, belinostat, CUDC-101, panobinostat, resminostat, RGFP966, rocilinostat, SB939, and tubacin. We also added mocetinostat and GSK J4 to this validation, since they were the two compounds showing reprogramming effect. Each inhibitor was screened at nine concentrations and four replicates for each dose. Only AR-42, mocetinostat, and GSK-J4 induced reprogramming in this second screening. We did not observe any sensitization for the other compounds and had to consider the low dose reprogramming as a measurement artifact, probably due to plate effect.

Three-day pretreatment resulted in the successful reprogramming of Oci-Ly-19, Riva-I, and Su-Dhl-4 cells, mainly by HDAC inhibitors. As most of the inhibitors were more effective after a 3-day pretreatment, we hypothesized that an even longer pretreatment time might increase the reprogramming effect. Hence, we decided to design our main screening assay with a 9-day pretreatment period and we extended our drug collection to 60 inhibitors.

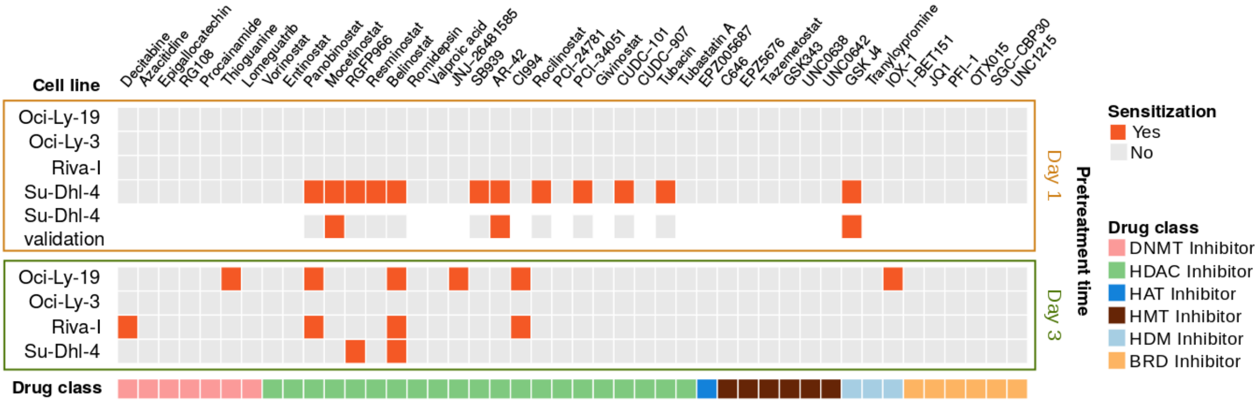


***Supplementary Figure S3:*** *Reprogramming screening hits. Reprogramming scores above a threshold of 30% (see Material and Methods) and whose dose-response curve passed visual inspection are considered as hits and marked in orange. Su-Dhl-4 validation row shows the results of the low dose effect assay.*

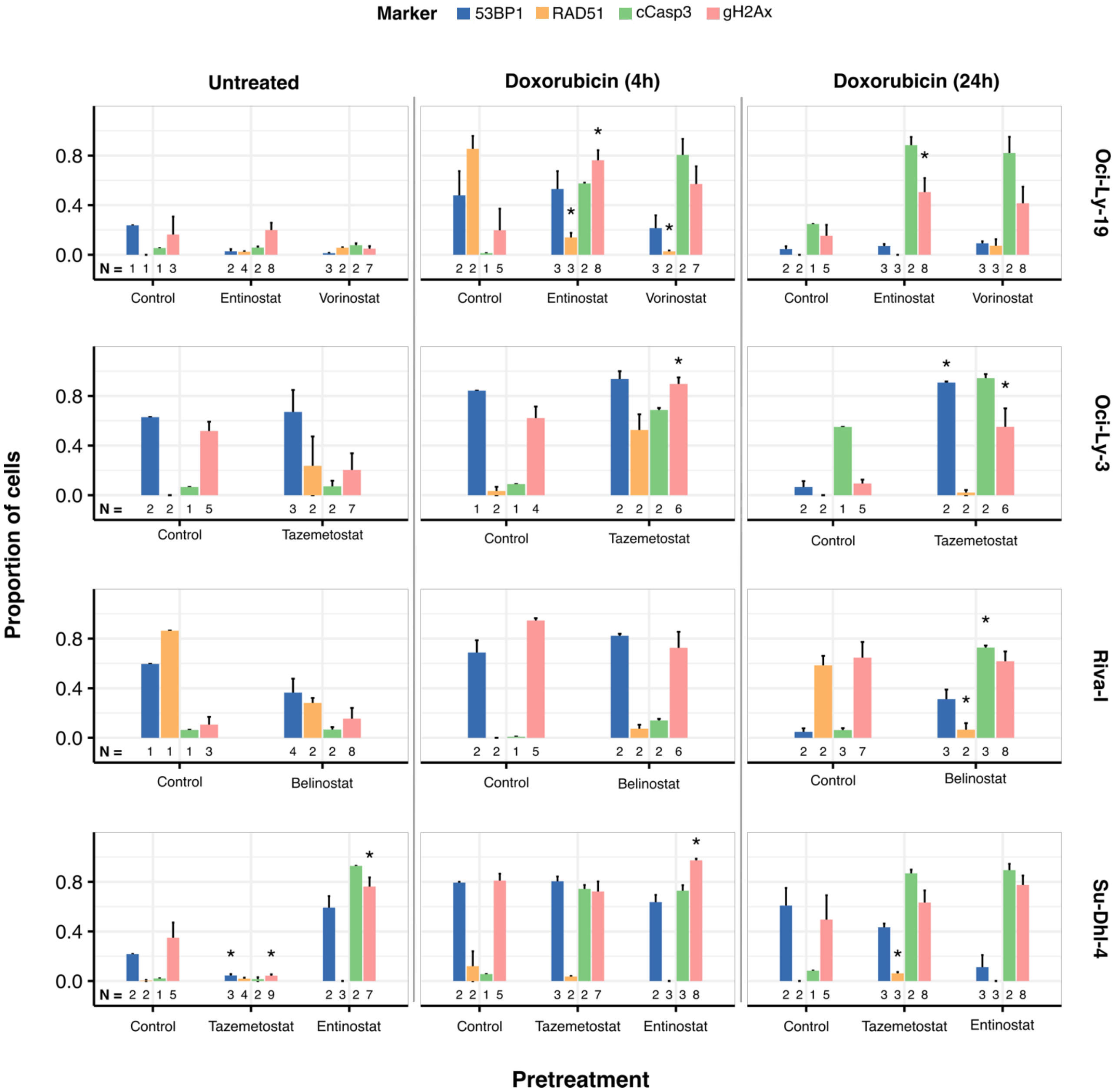


***Supplementary Figure S4:*** *Effect of doxorubicin treatment after epigenetic pretreatment on homologous recombination and DNA damage quantified by immunofluorescence assay. The markers 53BP1, RAD51, cCasp3, and gH2Ax represent non-homologous recombination DNA repair activation, homologous recombination DNA repair, apoptosis, and double strand DNA breaks respectively. The bar plots show doxorubicin effects on the protein expression on treated (marked with the corresponding inhibitor) and treatment-naive cells (Control), as well as epigenetic inhibitor effects in the absence of doxorubicin (left column marked as Untreated). Asterisks represent measurements significantly different from their respective Control in the cases where the number of images available (N) was sufficient for statistical test.*

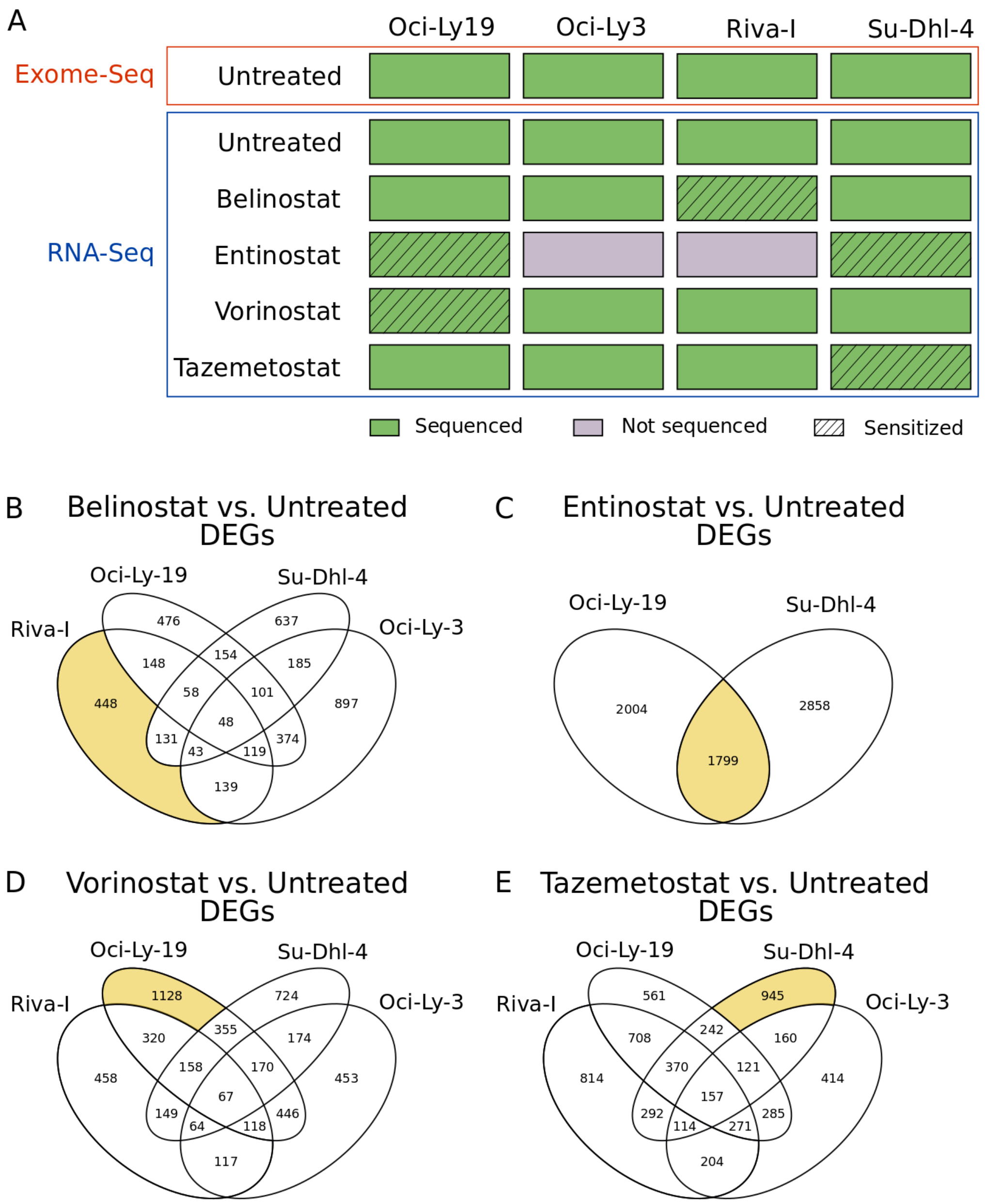


***Supplementary Figure S5:*** *Overview of sequencing data and selection of DEGs. (A) Summary of the sequencing experiments carried out in the study. Oci-Ly-3 and Riva-I cells treated with entinostat where not sequenced because the selected dose of entinostat was too toxic for these cell lines. (B-E) Venn diagrams showing the amount of DEGs induced by each epigenetic inhibitor. The portions of the diagram in yellow represent the DEGs of interested because they are found only in the sensitized combinations (B, D, and E) or are shared among the sensitized combinations (C).*

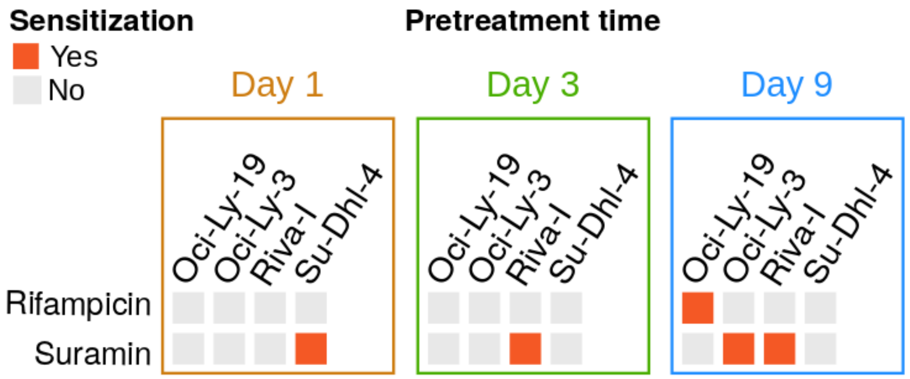


***Supplementary Figure S6:*** *Sensitizing effect of CD20-transport associated compounds.*
